# First genome-based characterization of *Listeria monocytogenes* in Costa Rica

**DOI:** 10.1101/2023.06.23.543262

**Authors:** María Giralt-Zúñiga, Mauricio Redondo-Solano, Alexandra Moura, Nathalie Tessaud-Rita, Hélène Bracq-Dieye, Guillaume Vales, Pierre Thouvenot, Alexandre Leclercq, Carolina Chaves-Ulate, Kattia Núñez-Montero, Rossy Guillén-Watson, Olga Rivas-Solano, Grettel Chanto-Chacón, Francisco Duarte-Martínez, Vanessa Soto-Blanco, Javier Pizarro-Cerdá, Marc Lecuit

## Abstract

Genomic data on the foodborne pathogen *Listeria monocytogenes* from Central America are scarse. We analysed 92 isolates collected in Costa Rica over a decade from different regions, compared them to publicly available genomes and identified unnoticed outbreaks. This study calls for mandatory reporting of listeriosis to improve pathogen surveillance.

---

*Listeria monocytogenes* (*Lm*) is a ubiquitous Gram-positive bacterium and one of the most deadly foodborne human pathogens. *Lm* is the causative agent of listeriosis, a severe infection with high hospitalization and mortality rates in at-risk populations (the elderly and immunocompromised, pregnant women and newborns) (*1*).

The *Lm* population is heterogeneous and can be classified into lineages (*2*), genoserogroups (*3*), clonal complexes (CCs) and sequence types (STs) as defined by multilocus sequence typing (MLST) (*4*), and sublineages (SLs) and cgMLST types (CTs) as defined by core genome MLST (cgMLST) (*5*). Major CCs and SLs are global (*5–7*) and can be heterogeneous in terms of virulence (*8,9*), with those from serogroup IVb (lineage I) often causing the most severe infections (*9*).

Pathogen surveillance using whole-genome sequencing (WGS) provides an unprecedented resolution for identifying clusters of cases and sources of contamination, as well as predicting the virulence and antimicrobial resistance profiles of strains, useful for risk assessment (*5,10*).

Previous studies have confirmed the presence of *Lm* in various foods in Costa Rica, with contamination levels ranging from 5% to 20% in processed meat products and fresh cheese, respectively (*11,12*). However, because listeriosis is not yet a notifiable disease in Costa Rica, data are scarce and the impact of the disease is unknown.

Here we used WGS to characterize 92 *Lm* recovered over an 11-year period from clinical (*n*=16), food (*n*=64), and production environment (*n*=9) samples in Costa Rica (**Appendix**), to understand the diversity and potential risk of circulating strains. Isolates for which location data were available were from urban areas (metropolitan area, including the capital city San José) and from rural areas where fresh cheese production is prevalent (Alajuela, Naranjo, San Ramón, Vara Blanca, Upala, and Turrialba, the latter accounting for 70% of fresh cheese produced in Costa Rica) (**Appendix**).

Isolates from lineage I (*n*=88, 95%) and lineage II (*n*=4, 5%) were unevenly distributed into 12 different sublineages (SL) and clonal complexes (CC) (**Figure 1, Appendix**). These included a new sublineage (designated SL1079, new MLST singleton ST1079) within lineage I with an atypical genoserogroup IIb profile (*lmo0737*^+^/*lmo1118*^-^/ORF2110^-^/ORF2819^+^/*prs*^+^, designated IIb-v1), represented by an isolate from shrimp. PCR and WGS confirmed the presence of *lmo0737* and flanking genes (*lmo0733*-*lmo0739*), typically present in *Lm* lineage II isolates (serogroups IIa and IIc) but only occasionally present in lineage I (serogroup IVb-v1) (*3,13*). Surprisingly, the most common sublineages in both clinical and food-associated sources were SL2/CC2 (*n*=61, 66%) and SL3/CC3 (*n*=13, 14%), accounting for 80% of the isolates. SL2/CC2 (serogroup IVb) and SL3/CC3 (serogroup IIb) isolates are found worldwide (*5,6*), but are rarely the most prevalent genotypes, although previous studies have shown a significant association of SL2/CC2 with severe infections (*9*). Available data from other countries in Central America confirmed the overrepresentation of SL2/CC2 and SL3/CC3 in Costa Rica (**Figure 1**). Whether this is due to the country’s geographical location, climatic peculiarities, commercial trends and/or natural reservoirs remains to be determined.

**Figure 1.**
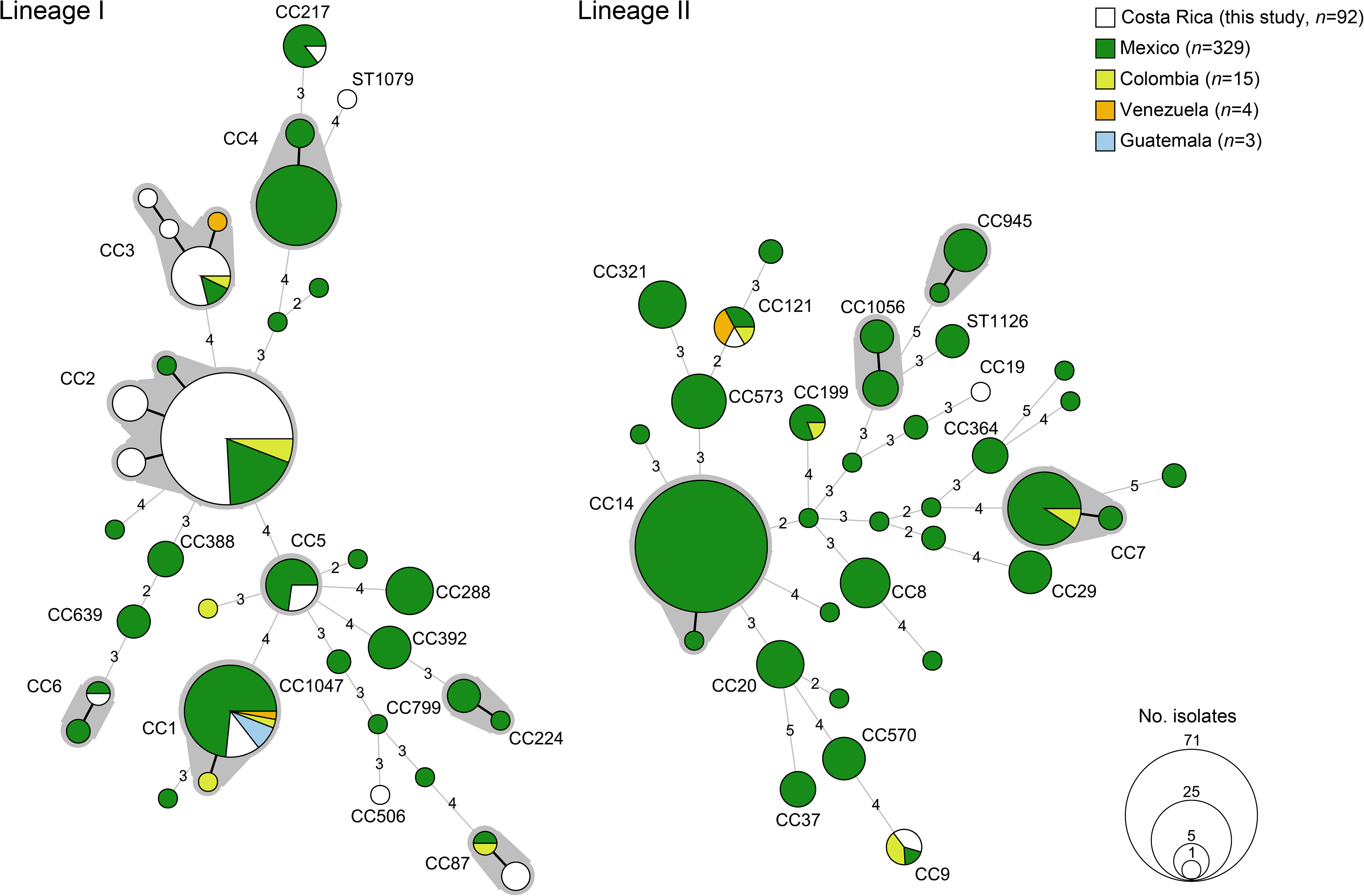
Minimum spanning tree of *L. monocytogenes* isolates from Costa Rica (*n*=92) based on MLST allelic profiles (7-locus scheme). Publicly available *L. monocytogenes* isolates (*n*=351) from neighboring countries in the Caribbean region were also included. Circles represent different profiles and sizes are proportional to the number of isolates within. Branch lengths are proportional to the allelic differences between the profiles which are indicated in the branches. For simplicity, allelic differences of 1 are omitted and represented by thicker branch lines. Clonal complexes with more than 1 profile are surrounded by gray shading and labelled if detected in this study or if they contain 5 or more isolates.

At the strain level, 48 cgMLST types (CTs) were identified, of which 44 (92%) were reported for the first time. 11 CTs (23%) included multiple isolates (cut-off of 7 allelic differences out of 1748 cgMLST loci (*5*)) and 4 CTs (8%) included up to 4 clinical cases (**Table 1, Appendix**). CgMLST type L1-SL2-ST2-CT2715 (*n*=8 isolates) accounted for 25% of clinical cases and spanned up to 9 years (**Table 1**).

**Table 1.** cgMLST types detected in this study comprising more than 2 isolates (*n*=11 out of 48; cut-off of 7 allelic differences).

| cgMLST type<br>(clonal complex, serogroup) | No.<br>isolates<br>(%, $N=92$) | No. clinical<br>isolates<br>(%, $n=16$) | No. non-<br>clinical<br>isolates (%,<br>$n=76$) | Food type | Isolation years | Genetic traits of<br>resistance |
| --- | --- | --- | --- | --- | --- | --- |
| *Lineage I* |  |  |  |  |  |  |
| L1-SL2-ST2-CT2715 (CC2, IVb) | 8 (9%) | 4 (25%) | 4 (5%) | Dairy, meat | 2009, 2011, 2013, 2016-2017 | *bcrABC*, *qacA*, LGI-2 |
| L1-SL2-ST2-CT6120 (CC2, IVb) | 10 (11%) | 2 (13%) | 8 (9%) | Dairy | 2010, 2013, 2016, 2018-2019 | *qacA*, LGI-2 |
| L1-SL2-ST2-CT2718 (CC2, IVb) | 5 (5%) | 1 (6%) | 4 (5%) | Dairy | 2016, 2019 | *qacA* |
| L1-SL2-ST1251-CT2780 (CC2, IVb) | 3 (3.3%) | 1 (6%) | 2 (3%) | Meat | 2015-2016, 2018 | *qacA* |
| L1-SL3-ST3-CT2730 (CC3, IIb) | 9 (10%) |  | 9 (10%) | Fish, meat | 2016 | *bcrABC*, SSI-1 |
| L1-SL2-ST2-CT6072 (CC2, IVb) | 5 (5%) |  | 5 (7%) | Dairy | 2019 | LGI-2 |
| L1-SL2-ST1627-CT6041 (CC2, IVb) | 5 (5%) |  | 5 (7%) | Dairy | 2018-2019 | LGI-2 |
| L1-SL87-ST847-CT65 (CC87, IIb) | 2 (2.2%) |  | 2 (3%) | Meat | 2016, 2019 | - |
| L1-SL3-ST1262/ST2762-CT2781 (CC3, IIb) | 2 (2.2%) |  | 2 (3%) | Dairy | 2013 | SSI-1 |
| L1-SL5-ST5-CT2793 (CC5, IIb) | 2 (2%) |  | 2 (3%) | Fish, meat | 2016 | *bcrABC*, SSI-1 |
| L1-SL2-ST2-CT2762 (CC2, IVb) | 2 (2%) |  | 2 (3%) | Mushrooms | 2011 | LGI-2 |

Most clusters with human cases were associated with dairy products (**Table 1**). However, tracing to confirm the source of infection was not possible because most production is carried out by local farmers throughout the country, often without traceability.

Fresh cheese production is an important economic income in Costa Rica and previous studies have reported the regular presence of *Lm* in such products (*12*). Our results also show the presence of identical strains (cgMLST type L1-SL2-ST2-CT6072) along the same production line, from raw materials to the final product, suggesting inadequate sanitation favoring contamination (*14,15*). Indeed, *Lm* is problematic for the food industry as it can often survive and multiply under adverse environmental conditions (*16*). Here, 90% of the isolates carried at least one genetic element encoding for tolerance to disinfectants [*qacA* (*n=*47, 51%); *bcrABC*, (*n*=21, 23%); *ermC* (n=1; 1%)], survival to low pH and high salt concentrations [SSI-1 (*n*=19, 21%)], high pH and oxidative stress [SSI-2 (*n*=1, 1%)] and/or to metals [LGI-2 (*n=*48, 50%); LGI-3, (*n*=1, 1%)], which could make *Lm* elimination from production sites more difficult.

Strengthening WGS surveillance in Costa Rica would be of great value to producers, informing them of strain diversity and effective means of eradication, while also allowing authorities to detect outbreaks and trace the sources of contamination in order to reduce the burden of listeriosis.

This study provides the first insight into the diversity of *Lm* strains circulating in Central America and calls for mandatory reporting of listeriosis to improve pathogen surveillance.

## Acknowledgments

This work includes MLST profiles publicly available on BIGSdb-*Listeria*. We thank the submitters for depositing their data in public databases and Institut Pasteur teams for the curation and maintenance of BIGSdb-Pasteur databases at https://bigsdb.pasteur.fr/. We also thank the CONAGEBIO from the Costa Rican Ministry of Environment and Energy (MINAE) for providing the permits to the access to biological material presented here (file CM-ITCR-003-2021).

## About the Author

Ms. Giralt-Zúñiga is a PhD student at the Molecular Microbiology department of the Institute for Biology, Humboldt-Universität zu Berlin, Germany. Her research interests are focused on infectious diseases and enteric pathogens.

## Appendix

### Materials and Methods

#### Bacterial isolation

The study included 92 *L. monocytogenes* isolates previously collected from different regions throughout Costa Rica (**Figure S1, Table S1**) and spanning eleven years (2009-2019). Clinical isolates (*n*=16) were obtained by the Institute for Research and Teaching in Nutrition and Health (INCIENSA) who receives clinical samples sent by hospitals across the country and by the University of Costa Rica (UCR). Isolates from food and food-production environments (*n*=76) were obtained by the National Laboratory of Veterinary Services (LANASEVE) of National Animal Health Service (SENASA), which performs microbiology analysis for the surveillance of food safety in products of animal origin for human consumption, the INCIENSA that also monitors the microbiological quality of food for human consumption, and by the UCR and the Instituto Tecnológico de Costa Rica (ITCR) in the scope of research projects and/or routine analyses for customers.

Bacterial isolation was performed as described previously, following either the *Bacteriological Analytical Manual* method for *Listeria* isolation (*1*), for isolates obtained by the UCR, ITCR and SENASA or the ISO 11290-1:2017 method (*2*), for isolates obtained by the INCIENSA.

Isolate identification was performed by proteomic analysis by Laser-Assisted Matrix and Flight Time Desorption/Ionization Mass Spectrometry (MALDI-TOF/MS), using the MicroFlex LT system with MBT library DB-5989 (Bruker Daltonics, Bremen, Germany) as previously described (*3*).

#### DNA extraction and genome sequencing

Isolates were cultured in Brain Heart Infusion Broth (BHI, Oxoid, Basingstoke UK) at 35 °C overnight before use. DNA extraction was performed with the DNA Easy Blood & Tissue Kit (QIAGEN, København Ø, Denmark-confirm), according to the instructions provided by the manufacturer. Qubit fluorometer (Thermo FisherScientific, Waltham, MA, USA) was used to evaluate DNA quantity and purity. Library preparation was performed with the Nextera XT DNA Sample Kit (Illumina, San Diego, CA, USA), and DNA sequencing was carried out on a NextSeq 500 platform (Illumina) using 2×150 bp paired-end runs. Reads were trimmed using fqCleanER v.21.10 (https://gitlab.pasteur.fr/GIPhy/fqCleanER), and assemblies were obtained using SPAdes v.3.14.0 (*4*) and polished with Pilon v.1.23 (*5*).

#### *In silico* molecular typing

PCR-serogroups (*6*), multilocus sequence types (MLST) (7), core genome MLST (cgMLST) (*8*), and virulence and resistance profiles (*8–14*) were extracted from draft assemblies using BIGSdb-*Lm* (*8,15*) and BLASTN algorithm, as described before (*8*).

Minimum spanning trees were obtained from MLST and cgMLST profiles using BioNumerics v.7.6 (Applied-Maths, Sint-Martens-Latem, Belgium). MLST analyses also included 351 publicly available *L. monocytogenes* profiles from neighboring countries, obtained from BIGSdb*-Lm* (http://bigsdb.pasteur.fr/listeria/; accessed on 16 February 2023). cgMLST-based dendrograms were built in BioNumerics v.7.6.3 (Applied Maths, Sint-Martens-Latem, Belgium) using categorical differences and the single-linkage clustering method, and visualezed with iTOL v.5 (*16*).

#### Data availability

Sequence data was made publicly available in NCBI/EBI/DDJJ databases (BioProject no. PRJEB20026).

---

**Appendix Figure 1.**
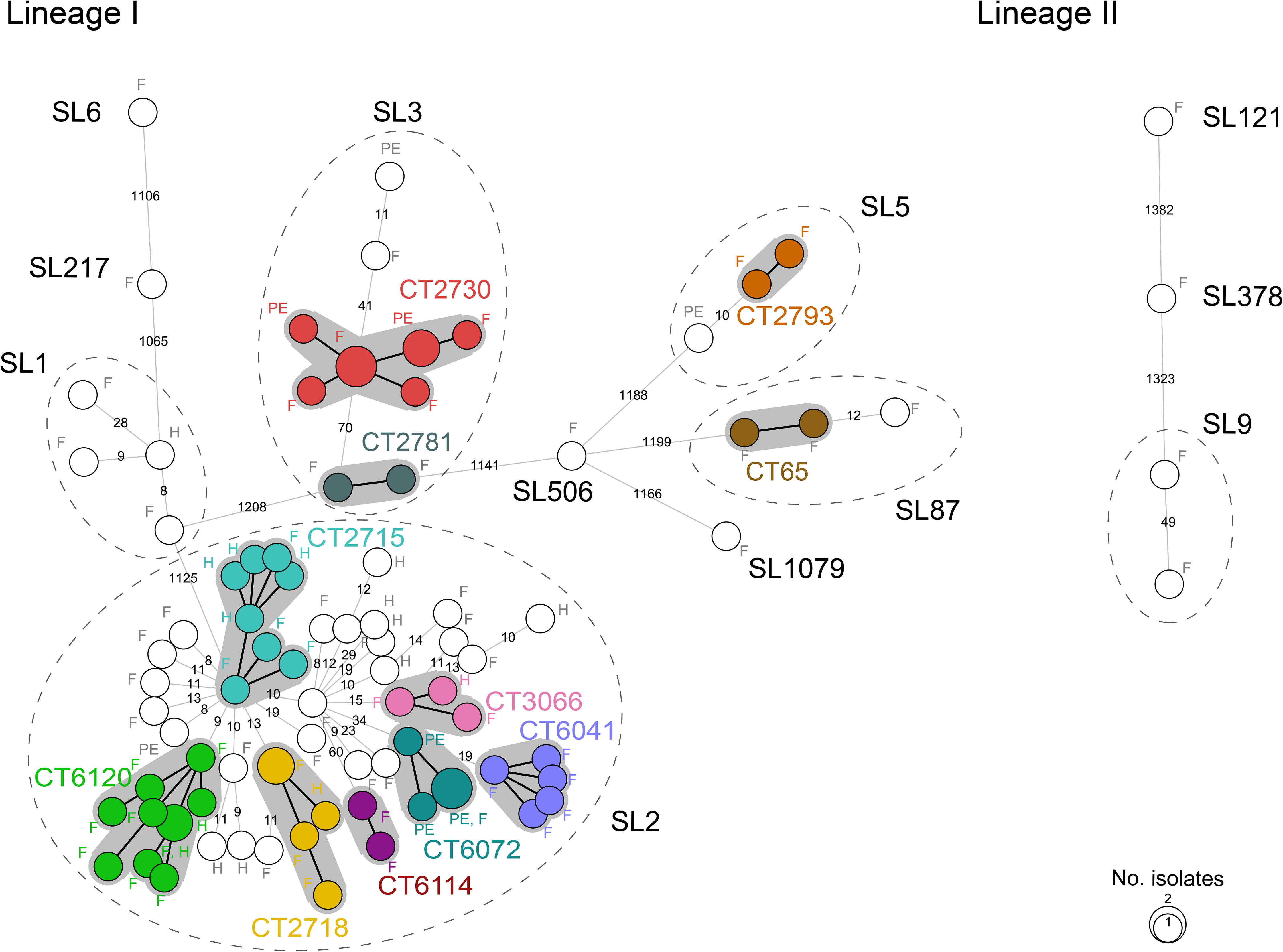
Minimum spanning tree of *L. monocytogenes* isolates from Costa Rica (*N*=92) based on cgMLST allelic profiles (1748-locus scheme). Circles represent different profiles and sizes are proportional to the number of isolates within. Labels next to circles indicate the source of isolates (H, human; F, food; PE, production environment). Branch lengths are proportional (in logarithmic scale) to allelic differences between profiles which are also indicated in the branches. For simplicity, allelic differences of 7 and below are omitted and represented by thicker branch lines. Clusters with more than 1 profile are highlighted in colors, labelled with corresponding cgMLST type, and delimitated by gray shadows. Dashed ellipses delimitate sublineages with more than 1 isolate and labelled with corresponding sublineage.

**Appendix Figure 2.**
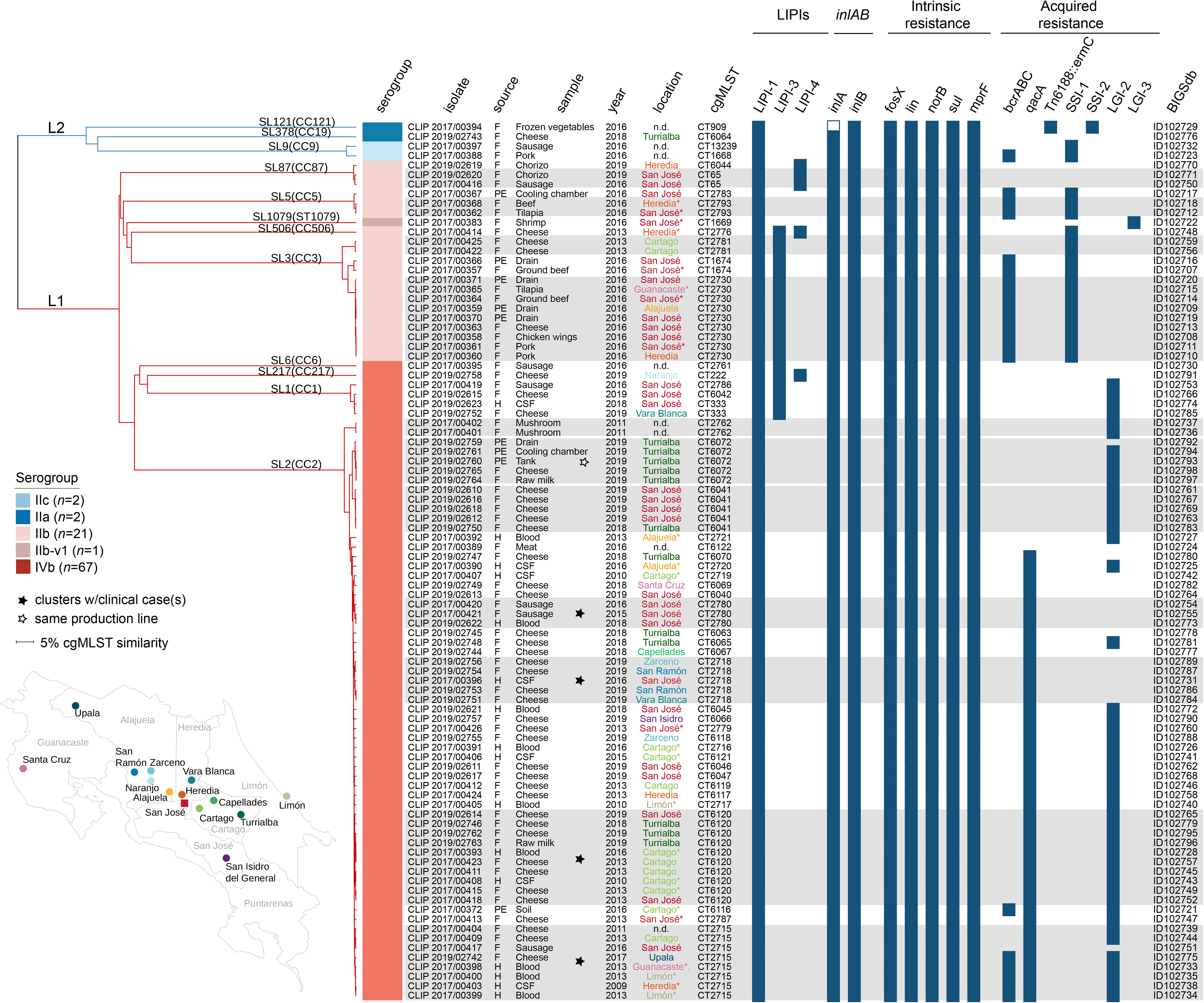
Single linkage dendrogram of *L. monocytogenes* isolates from Costa Rica (*N*=92) based on cgMLST allelic profiles (1748-locus scheme). Branches are colored according to lineages (lineage 1, red; lineage 2, blue) and labeled according to lineages and sublineages (SL) and clonal complexes (CC). Information on isolates’ name, serogroup (colored as in the key panel), source (H, human; F, food, PE, production environment), sample type, cgMLST type, year of isolation, location (colored as displayed on the map; asterisks indicate information available only at the level of provinces, which are labelled in gray; n.d., not determined) and BIGSdb ids are provided in the columns. Clusters of isolates with 7 or less allelic differences (out of 1748 cgMLST loci) are highlighted in gray boxes. Black stars highlight clusters of isolates containing human isolates. The presence of selected virulence and resistance genetic traits in each isolate is represented by squared dark blue boxes and empty boxes denote genes with premature stop codons.

**Appendix Table 1.** Isolate metadata and genome metrics of *Listeria monocytogenes* sequenced in this study (*N=*92).

| isolate | no. bases<br>after<br>filtering | coverage | no.<br>contigs | total length<br>(bp) | N50 (bp) | %GC | %<br>cgMLST<br>loci tagged | source<br>type | sample<br>type | Geographic<br>location | year of<br>isolation | serogroup | lineage | clonal<br>complex<br>(MLST) | sublineage<br>(cgMLST) | cgMLST<br>type<br>(cgMLST) | BIGSdb id |
| --- | --- | --- | --- | --- | --- | --- | --- | --- | --- | --- | --- | --- | --- | --- | --- | --- | --- |
| CLIP 2017/00419 | 2.40E+08 | 83 | 38 | 3.01E+06 | 1.94E+05 | 37.8 | 99.7 | F | Sausage | San José | 2016 | IVb | I | CC1 | SL1 | CT2786 | ID102753 |
| CLIP 2019/02752 | 3.82E+08 | 132 | 30 | 2.99E+06 | 2.62E+05 | 37.82 | 99.8 | F | Cheese | Vara Blanca | 2019 | IVb | I | CC1 | SL1 | CT333 | ID102785 |
| CLIP 2019/02623 | 4.20E+08 | 145 | 43 | 3.04E+06 | 1.92E+05 | 37.8 | 99.9 | H | CSF | San José | 2018 | IVb | I | CC1 | SL1 | CT333 | ID102774 |
| CLIP 2019/02615 | 5.11E+08 | 176 | 38 | 3.02E+06 | 2.55E+05 | 37.81 | 99.9 | F | Cheese | San José | 2019 | IVb | I | CC1 | SL1 | CT6042 | ID102766 |
| CLIP 2017/00404 | 3.05E+08 | 105 | 58 | 3.06E+06 | 1.43E+05 | 37.82 | 99.9 | F | Cheese | n.d. | 2011 | IVb | I | CC2 | SL2 | CT2715 | ID102739 |
| CLIP 2017/00409 | 2.52E+08 | 87 | 110 | 3.05E+06 | 7.45E+04 | 37.85 | 99.3 | F | Cheese | Cartago | 2013 | IVb | I | CC2 | SL2 | CT2715 | ID102744 |
| CLIP 2017/00417 | 5.93E+08 | 205 | 49 | 3.01E+06 | 1.39E+05 | 37.88 | 99.8 | F | Sausage | San José | 2016 | IVb | I | CC2 | SL2 | CT2715 | ID102751 |
| CLIP 2019/02742 | 5.20E+08 | 179 | 47 | 3.17E+06 | 1.98E+05 | 37.7 | 99.8 | F | Cheese | Upala | 2017 | IVb | I | CC2 | SL2 | CT2715 | ID102775 |
| CLIP 2017/00398 | 4.34E+08 | 150 | 58 | 3.17E+06 | 1.50E+05 | 37.69 | 99.8 | H | Blood | Guanacaste* | 2013 | IVb | I | CC2 | SL2 | CT2715 | ID102733 |
| CLIP 2017/00399 | 4.18E+08 | 144 | 51 | 3.16E+06 | 1.32E+05 | 37.69 | 99.8 | H | Blood | Limón* | 2013 | IVb | I | CC2 | SL2 | CT2715 | ID102734 |
| CLIP 2017/00400 | 3.59E+08 | 124 | 40 | 3.10E+06 | 2.23E+05 | 37.75 | 99.8 | H | Blood | Limón* | 2013 | IVb | I | CC2 | SL2 | CT2715 | ID102735 |
| CLIP 2017/00403 | 5.40E+08 | 186 | 52 | 3.17E+06 | 1.39E+05 | 37.69 | 99.8 | H | CSF | Heredia* | 2009 | IVb | I | CC2 | SL2 | CT2715 | ID102738 |
| CLIP 2017/00391 | 4.72E+08 | 163 | 51 | 3.04E+06 | 1.61E+05 | 37.84 | 99.9 | H | Blood | Cartago* | 2016 | IVb | I | CC2 | SL2 | CT2716 | ID102726 |
| CLIP 2017/00405 | 4.46E+08 | 154 | 38 | 3.05E+06 | 1.39E+05 | 37.83 | 99.9 | H | Blood | Limón* | 2010 | IVb | I | CC2 | SL2 | CT2717 | ID102740 |
| CLIP 2019/02751 | 4.38E+08 | 151 | 34 | 3.06E+06 | 2.73E+05 | 37.84 | 99.9 | F | Cheese | Vara Blanca | 2019 | IVb | I | CC2 | SL2 | CT2718 | ID102784 |
| CLIP 2019/02753 | 3.36E+08 | 116 | 29 | 3.06E+06 | 2.72E+05 | 37.84 | 99.9 | F | Cheese | San Ramón | 2019 | IVb | I | CC2 | SL2 | CT2718 | ID102786 |
| CLIP 2019/02754 | 3.03E+08 | 105 | 26 | 2.97E+06 | 3.33E+05 | 37.9 | 99.9 | F | Cheese | San Ramón | 2019 | IVb | I | CC2 | SL2 | CT2718 | ID102787 |
| CLIP 2019/02756 | 2.54E+08 | 87 | 58 | 3.01E+06 | 1.44E+05 | 37.88 | 99.8 | F | Cheese | Zarcero | 2019 | IVb | I | CC2 | SL2 | CT2718 | ID102789 |
| CLIP 2017/00396 | 5.98E+08 | 206 | 51 | 3.01E+06 | 1.41E+05 | 37.88 | 99.9 | H | CSF | San José | 2016 | IVb | I | CC2 | SL2 | CT2718 | ID102731 |
| CLIP 2017/00407 | 4.45E+08 | 153 | 58 | 3.06E+06 | 1.52E+05 | 37.83 | 99.9 | H | CSF | Cartago* | 2010 | IVb | I | CC2 | SL2 | CT2719 | ID102742 |
| CLIP 2017/00390 | 4.06E+08 | 140 | 42 | 3.04E+06 | 2.02E+05 | 37.81 | 99.9 | H | CSF | Alajuela* | 2016 | IVb | I | CC2 | SL2 | CT2720 | ID102725 |
| CLIP 2017/00392 | 5.23E+08 | 180 | 59 | 3.03E+06 | 1.52E+05 | 37.81 | 99.8 | H | Blood | Alajuela* | 2013 | IVb | I | CC2 | SL2 | CT2721 | ID102727 |
| CLIP 2017/00401 | 3.46E+08 | 119 | 38 | 2.96E+06 | 1.50E+05 | 37.85 | 99.8 | F | Mushroom | n.d. | 2011 | IVb | I | CC2 | SL2 | CT2762 | ID102736 |
| CLIP 2017/00402 | 2.98E+08 | 103 | 42 | 2.99E+06 | 1.38E+05 | 37.81 | 99.8 | F | Mushroom | n.d. | 2011 | IVb | I | CC2 | SL2 | CT2762 | ID102737 |
| CLIP 2017/00426 | 2.63E+08 | 91 | 55 | 3.04E+06 | 1.87E+05 | 37.84 | 99.9 | F | Cheese | San José* | 2013 | IVb | I | CC2 | SL2 | CT2779 | ID102760 |
| CLIP 2017/00420 | 1.59E+08 | 55 | 87 | 3.05E+06 | 9.85E+04 | 37.83 | 99.9 | F | Sausage | San José | 2016 | IVb | I | CC2 | SL2 | CT2780 | ID102754 |
| CLIP 2017/00421 | 2.61E+08 | 90 | 59 | 3.05E+06 | 1.53E+05 | 37.83 | 100 | F | Sausage | San José | 2015 | IVb | I | CC2 | SL2 | CT2780 | ID102755 |
| CLIP 2019/02622 | 3.75E+08 | 129 | 48 | 3.01E+06 | 2.38E+05 | 37.86 | 99.9 | H | Blood | San José | 2018 | IVb | I | CC2 | SL2 | CT2780 | ID102773 |
| CLIP 2017/00413 | 5.63E+08 | 194 | 61 | 3.08E+06 | 1.29E+05 | 37.79 | 99.9 | F | Cheese | San José* | 2013 | IVb | I | CC2 | SL2 | CT2787 | ID102747 |
| CLIP 2019/02613 | 4.08E+08 | 141 | 41 | 3.00E+06 | 1.73E+05 | 37.86 | 99.9 | F | Cheese | San José | 2019 | IVb | I | CC2 | SL2 | CT6040 | ID102764 |
| CLIP 2019/02610 | 4.27E+08 | 147 | 67 | 3.05E+06 | 1.45E+05 | 37.79 | 99.9 | F | Cheese | San José | 2019 | IVb | I | CC2 | SL2 | CT6041 | ID102761 |
| CLIP 2019/02612 | 3.79E+08 | 131 | 62 | 3.11E+06 | 1.61E+05 | 37.86 | 99.9 | F | Cheese | San José | 2019 | IVb | I | CC2 | SL2 | CT6041 | ID102763 |
| CLIP 2019/02616 | 3.38E+08 | 117 | 65 | 3.05E+06 | 1.11E+05 | 37.79 | 99.8 | F | Cheese | San José | 2019 | IVb | I | CC2 | SL2 | CT6041 | ID102767 |
| CLIP 2019/02618 | 3.65E+08 | 126 | 82 | 3.05E+06 | 1.10E+05 | 37.79 | 99.9 | F | Cheese | San José | 2019 | IVb | I | CC2 | SL2 | CT6041 | ID102769 |
| CLIP 2019/02750 | 4.94E+08 | 170 | 50 | 3.05E+06 | 1.98E+05 | 37.79 | 99.9 | F | Cheese | Turrialba | 2018 | IVb | I | CC2 | SL2 | CT6041 | ID102783 |
| CLIP 2019/02621 | 5.80E+08 | 200 | 51 | 3.08E+06 | 1.61E+05 | 37.84 | 99.9 | H | Blood | San José | 2018 | IVb | I | CC2 | SL2 | CT6045 | ID102772 |
| CLIP 2019/02611 | 3.92E+08 | 135 | 76 | 3.09E+06 | 1.98E+05 | 37.8 | 99.9 | F | Cheese | San José | 2019 | IVb | I | CC2 | SL2 | CT6046 | ID102762 |
| CLIP 2019/02617 | 3.55E+08 | 122 | 65 | 3.03E+06 | 1.26E+05 | 37.84 | 99.9 | F | Cheese | San José | 2019 | IVb | I | CC2 | SL2 | CT6047 | ID102768 |
| CLIP 2019/02745 | 3.90E+08 | 135 | 35 | 2.97E+06 | 2.40E+05 | 37.91 | 99.9 | F | Cheese | Turrialba | 2018 | IVb | I | CC2 | SL2 | CT6063 | ID102778 |
| CLIP 2019/02748 | 4.82E+08 | 166 | 43 | 3.04E+06 | 2.71E+05 | 37.85 | 99.9 | F | Cheese | Turrialba | 2018 | IVb | I | CC2 | SL2 | CT6065 | ID102781 |
| CLIP 2019/02757 | 3.04E+08 | 105 | 68 | 3.10E+06 | 2.39E+05 | 37.79 | 99.7 | F | Cheese | San Isidro | 2019 | IVb | I | CC2 | SL2 | CT6066 | ID102790 |
| CLIP 2019/02744 | 4.94E+08 | 170 | 51 | 3.04E+06 | 2.38E+05 | 37.85 | 99.8 | F | Cheese | Capellades | 2018 | IVb | I | CC2 | SL2 | CT6067 | ID102777 |
| CLIP 2019/02749 | 4.85E+08 | 167 | 40 | 3.04E+06 | 1.59E+05 | 37.85 | 99.9 | F | Cheese | Santa Cruz | 2018 | IVb | I | CC2 | SL2 | CT6069 | ID102782 |
| CLIP 2019/02747 | 4.08E+08 | 141 | 74 | 3.07E+06 | 1.98E+05 | 37.83 | 99.9 | F | Cheese | Turrialba | 2018 | IVb | I | CC2 | SL2 | CT6070 | ID102780 |
| CLIP 2019/02764 | 7.56E+08 | 261 | 84 | 3.16E+06 | 3.21E+05 | 37.76 | 99.8 | F | Raw milk | Turrialba | 2019 | IVb | I | CC2 | SL2 | CT6072 | ID102797 |
| CLIP 2019/02765 | 7.81E+08 | 269 | 59 | 3.12E+06 | 2.06E+05 | 37.79 | 99.9 | F | Cheese | Turrialba | 2019 | IVb | I | CC2 | SL2 | CT6072 | ID102798 |
| CLIP 2019/02759 | 1.24E+09 | 426 | 120 | 3.11E+06 | 3.60E+05 | 38 | 99.9 | PE | Drain | Turrialba | 2019 | IVb | I | CC2 | SL2 | CT6072 | ID102792 |
| CLIP 2019/02760 | 7.47E+08 | 258 | 47 | 3.12E+06 | 3.60E+05 | 37.77 | 99.9 | PE | Tank | Turrialba | 2019 | IVb | I | CC2 | SL2 | CT6072 | ID102793 |
| CLIP 2019/02761 | 7.06E+08 | 243 | 62 | 3.15E+06 | 3.63E+05 | 37.81 | 99.9 | PE | Cooling chamber | Turrialba | 2019 | IVb | I | CC2 | SL2 | CT6072 | ID102794 |
| CLIP 2017/00372 | 1.76E+08 | 61 | 90 | 3.12E+06 | 9.53E+04 | 37.69 | 99.9 | PE | Soil | Cartago* | 2016 | IVb | I | CC2 | SL2 | CT6116 | ID102721 |
| CLIP 2017/00424 | 5.53E+08 | 191 | 32 | 3.04E+06 | 1.98E+05 | 37.81 | 99.9 | F | Cheese | Heredia | 2013 | IVb | I | CC2 | SL2 | CT6117 | ID102758 |
| CLIP 2019/02755 | 3.66E+08 | 126 | 37 | 3.04E+06 | 3.55E+05 | 37.84 | 99.9 | F | Cheese | Zarcero | 2019 | IVb | I | CC2 | SL2 | CT6118 | ID102788 |
| CLIP 2017/00412 | 2.08E+08 | 72 | 60 | 3.06E+06 | 1.35E+05 | 37.8 | 99.8 | F | Cheese | Cartago | 2013 | IVb | I | CC2 | SL2 | CT6119 | ID102746 |
| CLIP 2017/00411 | 2.77E+08 | 96 | 57 | 3.08E+06 | 2.81E+05 | 37.8 | 99.8 | F | Cheese | Cartago | 2013 | IVb | I | CC2 | SL2 | CT6120 | ID102745 |
| CLIP 2017/00415 | 2.78E+08 | 96 | 51 | 3.04E+06 | 1.55E+05 | 37.84 | 99.9 | F | Cheese | Cartago* | 2013 | IVb | I | CC2 | SL2 | CT6120 | ID102749 |
| CLIP 2017/00418 | 2.74E+08 | 94 | 48 | 3.04E+06 | 1.50E+05 | 37.84 | 99.9 | F | Cheese | San José | 2013 | IVb | I | CC2 | SL2 | CT6120 | ID102752 |
| CLIP 2017/00423 | 2.62E+08 | 90 | 63 | 3.08E+06 | 1.39E+05 | 37.8 | 99.9 | F | Cheese | Cartago | 2013 | IVb | I | CC2 | SL2 | CT6120 | ID102757 |
| CLIP 2019/02614 | 2.43E+08 | 84 | 90 | 3.08E+06 | 9.05E+04 | 37.81 | 99.4 | F | Cheese | San José | 2019 | IVb | I | CC2 | SL2 | CT6120 | ID102765 |
| CLIP 2019/02746 | 3.45E+08 | 119 | 59 | 3.08E+06 | 1.61E+05 | 37.81 | 99.8 | F | Cheese | Turrialba | 2018 | IVb | I | CC2 | SL2 | CT6120 | ID102779 |
| CLIP 2019/02762 | 6.86E+08 | 236 | 475 | 3.27E+06 | 2.73E+05 | 39.67 | 99.8 | F | Cheese | Turrialba | 2019 | IVb | I | CC2 | SL2 | CT6120 | ID102795 |
| CLIP 2019/02763 | 9.18E+08 | 317 | 136 | 3.12E+06 | 3.58E+05 | 38.07 | 99.9 | F | Raw milk | Turrialba | 2019 | IVb | I | CC2 | SL2 | CT6120 | ID102796 |
| CLIP 2017/00393 | 4.76E+08 | 164 | 59 | 3.08E+06 | 1.50E+05 | 37.8 | 99.9 | H | Blood | Cartago* | 2016 | IVb | I | CC2 | SL2 | CT6120 | ID102728 |
| CLIP 2017/00408 | 4.09E+08 | 141 | 66 | 3.08E+06 | 1.36E+05 | 37.81 | 99.9 | H | CSF | Cartago* | 2010 | IVb | I | CC2 | SL2 | CT6120 | ID102743 |
| CLIP 2017/00406 | 6.63E+08 | 228 | 60 | 3.07E+06 | 1.38E+05 | 37.79 | 99.9 | H | CSF | Cartago* | 2015 | IVb | I | CC2 | SL2 | CT6121 | ID102741 |
| CLIP 2017/00389 | 3.43E+08 | 118 | 41 | 3.05E+06 | 1.39E+05 | 37.83 | 99.9 | F | Meat | n.d. | 2016 | IVb | I | CC2 | SL2 | CT6122 | ID102724 |
| CLIP 2019/02758 | 8.26E+08 | 285 | 31 | 2.91E+06 | 4.33E+05 | 37.91 | 99.8 | F | Cheese | Naranjo | 2019 | IVb | I | CC217 | SL217 | CT222 | ID102791 |
| CLIP 2017/00395 | 3.03E+08 | 104 | 41 | 3.00E+06 | 1.95E+05 | 37.79 | 99.9 | F | Sausage | n.d. | 2016 | IVb | I | CC6 | SL6 | CT2761 | ID102730 |
| CLIP 2017/00383 | 3.27E+08 | 113 | 59 | 3.03E+06 | 1.55E+05 | 37.82 | 99.9 | F | Shrimp | San José* | 2016 | IIb-v1 | I | ST1079 | SL1079 | CT1669 | ID102722 |
| CLIP 2017/00357 | 3.37E+08 | 116 | 53 | 3.14E+06 | 1.58E+05 | 37.75 | 99.8 | F | Ground beef | San José* | 2016 | IIb | I | CC3 | SL3 | CT1674 | ID102707 |
| CLIP 2017/00366 | 3.20E+08 | 110 | 51 | 3.10E+06 | 1.97E+05 | 37.79 | 99.9 | PE | Drain | San José | 2016 | IIb | I | CC3 | SL3 | CT1674 | ID102716 |
| CLIP 2017/00358 | 2.38E+08 | 82 | 45 | 3.05E+06 | 1.34E+05 | 37.81 | 99.8 | F | Chicken wings | San José | 2016 | IIb | I | CC3 | SL3 | CT2730 | ID102708 |
| CLIP 2017/00360 | 3.67E+08 | 127 | 43 | 3.05E+06 | 1.86E+05 | 37.81 | 99.8 | F | Pork | Heredia | 2016 | IIb | I | CC3 | SL3 | CT2730 | ID102710 |
| CLIP 2017/00361 | 5.29E+08 | 183 | 44 | 3.05E+06 | 1.87E+05 | 37.81 | 99.8 | F | Pork | San José* | 2016 | IIb | I | CC3 | SL3 | CT2730 | ID102711 |
| CLIP 2017/00363 | 4.02E+08 | 139 | 51 | 3.05E+06 | 1.42E+05 | 37.81 | 99.7 | F | Cheese | San José | 2016 | IIb | I | CC3 | SL3 | CT2730 | ID102713 |
| CLIP 2017/00364 | 4.97E+08 | 171 | 44 | 3.05E+06 | 1.69E+05 | 37.81 | 99.7 | F | Ground beef | San José* | 2016 | IIb | I | CC3 | SL3 | CT2730 | ID102714 |
| CLIP 2017/00365 | 2.15E+08 | 74 | 43 | 3.05E+06 | 1.97E+05 | 37.81 | 99.8 | F | Tilapia | Guanacaste* | 2016 | IIb | I | CC3 | SL3 | CT2730 | ID102715 |
| CLIP 2017/00359 | 3.82E+08 | 132 | 40 | 3.05E+06 | 3.71E+05 | 37.81 | 99.8 | PE | Drain | Alajuela | 2016 | IIb | I | CC3 | SL3 | CT2730 | ID102709 |
| CLIP 2017/00370 | 3.05E+08 | 105 | 40 | 3.05E+06 | 1.97E+05 | 37.81 | 99.8 | PE | Drain | San José | 2016 | IIb | I | CC3 | SL3 | CT2730 | ID102719 |
| CLIP 2017/00371 | 5.02E+08 | 173 | 42 | 3.05E+06 | 2.36E+05 | 37.81 | 99.7 | PE | Drain | San José | 2016 | IIb | I | CC3 | SL3 | CT2730 | ID102720 |
| CLIP 2017/00422 | 2.31E+08 | 80 | 39 | 3.04E+06 | 2.53E+05 | 37.92 | 99.7 | F | Cheese | Cartago | 2013 | IIb | I | CC3 | SL3 | CT2781 | ID102756 |
| CLIP 2017/00425 | 2.84E+08 | 98 | 39 | 3.04E+06 | 1.97E+05 | 37.93 | 99.7 | F | Cheese | Cartago | 2013 | IIb | I | CC3 | SL3 | CT2781 | ID102759 |
| CLIP 2017/00367 | 2.62E+08 | 90 | 79 | 3.11E+06 | 2.36E+05 | 37.79 | 99.9 | PE | Cooling chamber | San José | 2016 | IIb | I | CC5 | SL5 | CT2783 | ID102717 |
| CLIP 2017/00362 | 4.62E+08 | 159 | 74 | 3.11E+06 | 1.35E+05 | 37.77 | 99.9 | F | Tilapia | San José | 2016 | IIb | I | CC5 | SL5 | CT2793 | ID102712 |
| CLIP 2017/00368 | 5.08E+08 | 175 | 59 | 3.10E+06 | 1.53E+05 | 37.79 | 99.9 | F | Beef | Heredia* | 2016 | IIb | I | CC5 | SL5 | CT2793 | ID102718 |
| CLIP 2017/00414 | 2.13E+08 | 73 | 77 | 2.93E+06 | 1.20E+05 | 37.86 | 99.4 | F | Cheese | Heredia* | 2013 | IIb | I | CC506 | SL506 | CT2776 | ID102748 |
| CLIP 2019/02619 | 3.78E+08 | 130 | 39 | 2.97E+06 | 3.04E+05 | 37.88 | 99.9 | F | Chorizo | Heredia | 2019 | IIb | I | CC87 | SL87 | CT6044 | ID102770 |
| CLIP 2017/00416 | 2.36E+08 | 81 | 38 | 2.92E+06 | 1.99E+05 | 37.91 | 99.9 | F | Sausage | San José | 2016 | IIb | I | CC87 | SL87 | CT65 | ID102750 |
| CLIP 2019/02620 | 3.96E+08 | 137 | 39 | 2.97E+06 | 2.09E+05 | 37.88 | 99.9 | F | Chorizo | San José | 2019 | IIb | I | CC87 | SL87 | CT65 | ID102771 |
| CLIP 2017/00397 | 4.04E+08 | 139 | 28 | 3.09E+06 | 3.73E+05 | 37.87 | 100 | F | Sausage | n.d. | 2016 | IIc | II | CC9 | SL9 | CT13239 | ID102732 |
| CLIP 2017/00388 | 2.98E+08 | 103 | 44 | 3.04E+06 | 2.02E+05 | 37.8 | 100 | F | Pork | n.d. | 2016 | IIc | II | CC9 | SL9 | CT1668 | ID102723 |
| CLIP 2017/00394 | 3.41E+08 | 117 | 38 | 3.09E+06 | 2.01E+05 | 37.79 | 100 | F | Frozen vegetables | n.d. | 2016 | IIa | II | CC121 | SL121 | CT909 | ID102729 |
| CLIP 2019/02743 | 3.86E+08 | 133 | 18 | 2.83E+06 | 3.82E+05 | 37.93 | 99.9 | F | Cheese | Turrialba | 2018 | IIa | II | CC19 | SL378 | CT6064 | ID102776 |

\*Geographical information available only at the level of the province.

**Appendix Table 2.** Previously reported cgMLST types detected in this study (cut-off of 7 or less allelic differences out of 1748 cgMLST loci, Institut Pasteur scheme).

| cgMLST type<br>(clonal complex, serogroup) | source(s)<br>(this study) | source(s)<br>(other studies) | countries<br>(other studies) | source lab | NCBI/EBI/DDJJ<br>accession no. | references |
| --- | --- | --- | --- | --- | --- | --- |
| *Lineage I* |  |  |  |  |  |  |
| L1-SL217-ST217-CT222 (CC217, IVb) | F (dairy) | H,F (salad) | US | CDC | SRR1021894, SRR1027089 | (8) |
| L1-SL1-ST1-CT333 (CC1, IVb) | H,F (dairy) | H | US | CDC | SRR1043171, SRR7057542 | (8,17) |
| L1-SL87-ST847-CT65 (CC87, IIb) | F (meat) | F (avocado) | MX | FDA | SRR975360 | (8) |
| *Lineage II* |  |  |  |  |  |  |
| L2-SL121-ST121-CT909 (CC121, IIa) | F (frozen vegetables) | H,F,PE,FE,A | CL,DK,FR,DE,LV,NO,PL,NL,UK,US | KMAHVH,<br>ANSES, ECDC, IP,<br>UWLMO, UKHSA,<br>FDA, RKI, | ERR1304231, ERR1738638, ERR1738650,<br>ERR2522041, ERR2522263, ERR2522276,<br>ERR2522284, ERR2522292, ERR2522294,<br>ERR2522295, ERR2522297, ERR2522312,<br>ERR2522337, ERR2522338, ERR2522353,<br>ERR2522363, ERR2522371, ERR2522810,<br>ERR3040059, ERR3040061, ERR3040067,<br>ERR3040072, ERR3040073, ERR3040075,<br>ERR3040076, ERR3040078, ERR3040083,<br>ERR3040104, ERR4176463, ERR4176497,<br>ERR4176536, ERR4176606, ERR4176666,<br>ERR4176794, ERR4648214, SRR10753542,<br>SRR10753574, SRR10753599,<br>SRR10753600, SRR10753602,<br>SRR11004717, SRR13072765,<br>SRR14783257, SRR15245555,<br>SRR15245949, SRR20053048, SRR2040689,<br>SRR4052010, SRR4052014, SRR4052068,<br>SRR4052071, SRR4052072, SRR4052073,<br>SRR4052161, SRR4124934, SRR4124940,<br>SRR5133499, SRR5318940, SRR5526029,<br>SRR5526031, SRR5526035, SRR5526085,<br>SRR5526092, SRR5526094, SRR5526097,<br>SRR5526099, SRR5526101, SRR5526105,<br>SRR5526120, SRR5526126, SRR5526128,<br>SRR5526130, SRR5526135, SRR5526137,<br>SRR5526138, SRR5526142, SRR5526145,<br>SRR5526155, SRR5647016, SRR5647027,<br>SRR5647029, SRR5647030, SRR6966182,<br>SRR7403111, SRR7410624, SRR7429735,<br>SRR7429781, SRR7440567, SRR7440606,<br>SRR7440615, SRR7440618, SRR7440621,<br>SRR7440634, SRR7440649, SRR7440688,<br>SRR7440921, SRR7441075, SRR7441146, | (8,10,18,19) |

Abbreviations: Source - H, human; F, food; PE, food production environment; FE, farm environment; A, animal. Country - CL, Chile; DK, Denmark; FR, France; LV, Latvia; MX, Mexico; NO, Norway; PL, Poland; UK, United Kingdom; US, United States of America. Source lab - ANSES, Agence Nationale de Sécurité
Sanitaire de L'Alimentation, de L'Environnement et du Travail, FR; CDC, Centers for Disease Control and Prevention, US; ECDC, European Centre for Disease Prevention and Control; FDA, U.S. Food and Drug Administration, US; Institut Paster, FR; KMAHVH, Klinisk Mikrobiologisk Afdeling, DK; RKI, Robert Koch Institut, DE; UKHSA, UK Health Security Agency, UK; UWLMO, University of Warmia and Mazury in Olsztyn, PL.

